# VPS34-IN1 inhibits cap-mediated translation and synergizes with STING to drive type-I IFN expression in human plasmacytoid DCs

**DOI:** 10.1101/2024.06.17.599308

**Authors:** Paulo Antas, Mariana D. Machado, Fátima Leite-Pinheiro, Daniela Barros, Carlota Ramalhinho, Andreia Mendes, Beatriz H. Ferreira, Daniela Carvoeiro, Marisa Reverendo, Iola F. Duarte, Miwako Narita, Bing Su, Rafael J. Argüello, Beatrice Nal, Philippe Pierre, Catarina R. Almeida, Evelina Gatti

**Affiliations:** Institute of Biomedicine (iBiMED), Department of Medical Sciences, University of Aveiro, 3810-193 Aveiro, Portugal; Aix Marseille Université, CNRS, INSERM, CIML, 13288 Marseille cedex 9, France; Shanghai Institute of Immunology, Department of Microbiology and Immunology, Shanghai Jiao Tong University School of Medicine, Shanghai 200025, PR China; Niigata University, Faculty of Medicine, School of Health Sciences, Niigata 951-8518, Japan; CICECO, Aveiro Institute of Materials, Department of Chemistry, University of Aveiro, 3810-193 Aveiro, Portugal

**Keywords:** BDPCN, chemotherapy, CL307, immunotherapy, SGK3, Type-I interferon, VPS34

## Abstract

Inhibition of the phosphatidylinositol kinase vacuolar protein sorting 34 (VPS34) with the pharmacological compound VPS34-IN1 has a range of effects on the dynamics of endosomes. While VPS34 inhibition has been suggested as a potential therapeutic approach for treating certain cancers, our findings indicate that it has minimal cytotoxic effects on leukemic blastic plasmacytoid dendritic cell neoplasms (BPDCN). VPS34-IN1, however, interferes with plasmacytoid dendritic cells (pDCs) function by blocking the recruitment of serum and glucocorticoid-regulated kinase 3 (SGK3) to endosomes, which is shown to be necessary for Toll-like receptor 7 (TLR7) signaling. In a contrasting parallel, VPS34-IN1 triggers the activation of the stimulator of interferon genes (STING) and significantly enhances pDCs’ response to the STING agonist 2’3’-cyclic guanosine monophosphate-adenosine monophosphate (2’3’-cGAMP). This cooperative action with VPS34-IN1 leads to strongly increased expression of type-I interferons (IFNs), associated with an alteration of STING degradation and importantly, inhibition of cap-mediated mRNA translation. Inhibition of protein synthesis by VPS34-IN1 appears to be central to this synergy with STING activation, notably by compromising the expression of IFIT1/ISG56, a negative regulator of innate signaling. Thus, despite their limited toxicity towards different cancer lines, inhibitors targeting VPS34 and SGK3 may present promising compounds for controlling the expression of type-I IFNs in response to various microbial stimuli and pathological contexts.

**One-sentence summary:** Pharmacological inhibition of VPS34 affects multiple signaling pathways downstream of innate immunity receptors and consequently can inhibit or potentiate type-I Interferon induction according to the danger or microbial stimuli received by plasmacytoid DCs.

## Introduction

Endosomes and lysosomes, critical cellular organelles, play pivotal roles in orchestrating the uptake, degradation, and recycling of materials delivered via endocytosis or autophagy (*1*). These organelles are finely tuned systems, with specific subsets of endosomal compartments sharing a common repertoire of membrane proteins and governed by a phosphatidylinositol (PtdIns) lipid code that is instrumental in their identification, sorting, and dynamic regulation (*2*).

At the heart of this intricate PtdIns machinery lies vacuolar protein sorting 34 (VPS34), a member of the PtdIns lipid kinase family (*3*). VPS34 associates with the regulatory subunit VPS15 to form the sole class III PtdIns(3)P-kinase found in mammalian cells. These VPS34 complexes, comprising VPS34/VPS15 along with Beclin1, and either ATG14 or UVRAG, constitute complex I (CI) and complex II (CII), respectively (*3*). While CII predominantly governs endolysosomal sorting and autophagosome-lysosome fusion (*3,4,5*), CI plays a pivotal role in autophagosome biogenesis (*6*).

The accumulation of PtdIns(3)P, a consequence of VPS34 complex activity, serves as a beacon, attracting effector proteins equipped with PtdIns(3)P-interacting domains, such as FYVE, PX, and PROPPINS (*7*). These effectors are instrumental in regulating various cellular processes, including autophagy, endocytic sorting, phagocytosis, and cell signaling. Notably, the PX domain-containing serum and glucocorticoid-regulated kinase 3 (SGK3) is among these effectors, contributing to the intricate network of VPS34-mediated cellular activities (*3,8*).

Given the critical roles of these pathways in tumor development and growth and the availability of potent pharmacological inhibitors (*9*,*10*), the inhibition of VPS34 has emerged as a promising chemotherapeutic strategy for leukemia and breast cancer treatment (*11*,*12*). Inhibition of PtdIns(3)P production, also impinges on specialized functions in immune cells requiring endocytosis and autophagy. These functions encompass the recognition of microbial-associated molecular patterns (MAMPs), MHC-restricted antigen presentation, and cytokine production (*13*).

Due to the key role of these functions in immune responses, we initiated an investigation into the impact of VPS34 pharmacological inhibition on the leukemic blastic plasmacytoid dendritic cell neoplasm (BPDCN)-derived cell line CAL-1 (*14*), using VPS34-IN1, a highly selective and potent inhibitor of VPS34 (*8*). BPDCN represents a rare and aggressive hematologic malignancy characterized by the development of skin tumors, lymph node and splenic enlargement, as well as circulating leukemia or bone marrow infiltration. BPDCN originates from the precursors of type-I Interferon (IFN)-producing plasmacytoid dendritic cells (pDCs) and shares many phenotypic and functional properties with freshly isolated human blood pDCs (*15*, *16*). These properties include the expression of Toll-like receptor 7 (TLR7) and 9 (TLR9), which, upon transport to the endocytic pathway and interaction with nucleic acid (NA) ligands (*17*), trigger the expression of type-I-IFNs and other pro-inflammatory cytokines (*18*).

While VPS34-IN1 did not demonstrate marked cytotoxicity towards CAL-1 cells *in vitro*, it did curtail protein synthesis by inhibiting the mammalian target of rapamycin complex (mTORC)/ribosomal protein S6 kinase (S6K) pathway (*19*). In parallel, VPS34-IN1 was found to independently trigger the expression of type-I IFN mRNA, an effect attributed to the delayed degradation of the antiviral signaling adaptor stimulator of interferon genes (STING, STING1, TMEM173) (*20*). The combined effects of delayed STING degradation and reduced protein synthesis were found to synergize and strongly potentiate STING signaling upon VPS34-IN1 treatment. Conversely, although SGK3 inhibition by VPS34-IN1 had no effect on STING activity, VPS34-IN1 treatment or SGK3 targeted degradation displayed modulatory effects on TLR7 signaling (*8*). SGK3 was found to be crucial for type-I IFN mRNA induction in both primary and tumoral pDCs responding to TLR7 agonists, placing SGK3 downstream for TLR7 signaling.

In summary, our study unravels the intricate interplay between VPS34, SGK3 and STING in regulating innate receptor signaling, and by extension, the production of type-I IFN and pro-inflammatory cytokines in pDCs responding to various immunostimulatory cues. The pharmacological inhibition of these key players holds promise as a potential therapeutic strategy in the contexts of cancer and autoimmunity, with the potential to transform immunologically cold cancers into hot inflamed tumors (*21*).

## Results

### VPS34-IN1 impairs TLR7-dependent type-I IFN production in CAL-1 BPDCN

VPS34 complexes are known to control both autophagy and early endosome dynamics (graphical abstract Fig. EV1A) (*3*). We examined the conversion of ATG8/LC3-I into lipidated ATG8/LC3-II in the presence of the V-ATPase inhibitor bafilomycin A1. VPS34-IN1 treatment effectively prevented the time-dependent accumulation of LC3-II in CAL-1 cells (Fig. 1A). This finding strongly suggests the inhibition of macroautophagy flux in the pDC cell line, as suggested by previous observations (*22*). VPS34-IN1 treatment also led to a significant reduction in mitophagy, resulting in the accumulation of damaged mitochondria capable of producing reactive oxygen species (ROS), as detected by flow cytometry using the fluorescent MitoSOX dye (*23*) (Fig. 1B). We observed a marked perturbation in the distribution of the early endosome-associated FYVE domain-bearing EEA1 protein upon VPS34-IN1 treatment (Fig. 1C). This alteration in EEA1 localization in an organelle clustering aligns with our expectations of disrupted PtdIns(3)P generation in the early endocytic pathway following VPS34 inhibition (Fig. EV1A) (*24*).

**Figure 1.**
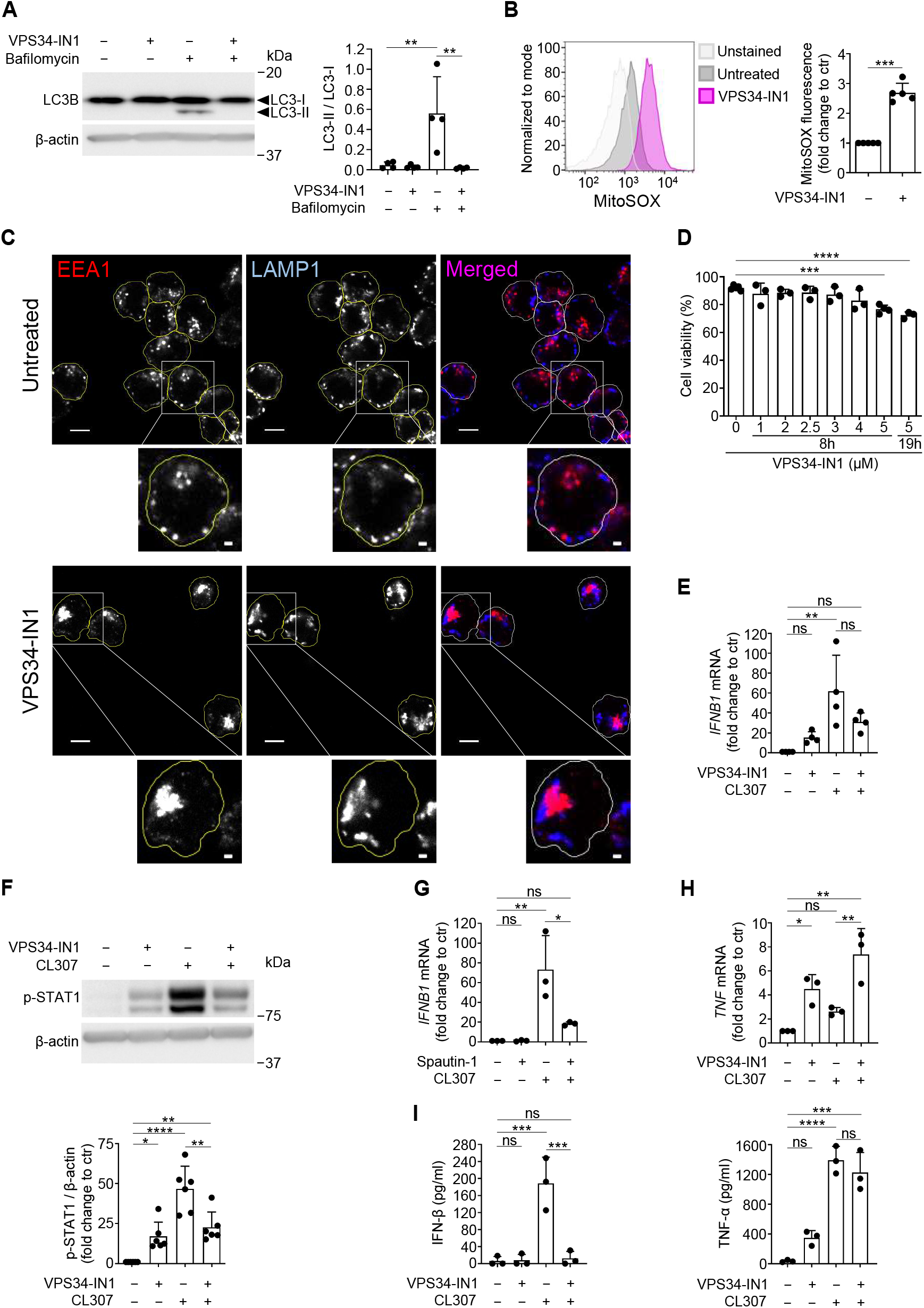
VPS34-IN1 impairs TLR7-mediated type-I IFN production in CAL-1. (**A**) LC3B lipidation was analyzed by determining the LC3-II/LC3-I ratio for CAL-1 cells treated with VPS34-IN1 for 8 h and bafilomycin for 4 h. LC3-I, nonlipidated form; LC3-II, lipidated form. (**B**) CAL-1 cells treated with VPS34-IN1 were stained with MitoSOX and analyzed by flow cytometry. The mean fluorescence intensity was compared to untreated cells (ctr). (**C**) Immunofluorescence confocal microscopy of VPS34-IN1-treated CAL-1 cells showing EEA1 (red) and LAMP1 (blue) distribution (n = 3). Scale bar, 5 μm or 1 μm in magnified areas. (**D**) Cell viability was determined for cells treated with varying concentration of VPS34-IN1 by propidium iodide exclusion and flow cytometry. (**E**-**I**) Cells were stimulated with CL307 and VPS34-IN1 or spautin-1. (**E, G, H**) Levels of *IFNB1* and *TNF* mRNA were normalized to housekeeping gene (*GAPDH*) and fold change determined in comparison with untreated cells (ctr). (**F**) Immunoblot for phosphorylated STAT1 (Tyr^701^). Levels of pSTAT1 relative to β-actin were compared against untreated cells (ctr). (**I**) Secretion of IFN-β and TNF-α was monitored by ELISA. (**A**, **B**, **D**-**I**) Bars show the mean ± SD and each dot shows data from one experiment. ns, non-significant results, *P < 0.05, **P < 0.01, ***P < 0.001, ****P < 0.0001, by one-way ANOVA with Šidák post hoc test (A), paired Student’s t test (B), one-way ANOVA with Dunnett post hoc test (D) and one-way ANOVA with Tukey post hoc test (E-I).

Given the efficacy of VPS34-IN1 in affecting known VPS34-dependent processes in CAL-1 cells, we conducted a dose-response assay to assess the cytotoxicity of VPS34-IN1 on this BPDCN line using propidium iodide (PI) staining (Fig. 1D). Our findings revealed a moderate level of cytotoxicity, with an approximately 20% reduction in cell viability observed at a dose of 5 µM of VPS34-IN1 over an 8-20 hours period. Notably, this cytotoxicity was only considered moderate when compared to the effects of other chemotherapeutic agents previously tested on this cell line (*25*). We next explored the response of CAL-1 cells to endosomal Toll-like receptor (TLR) agonists, specifically using CL307, a small compound known for its selective stimulation of TLR7 (*26*). Inhibition of LC3 lipidation, accumulation of ROS-producing mitochondria and alteration of early endosome distribution by VPS34-IN1 were all recapitulated in CL307 activated cells (Fig. EV2).

Upon treatment with VPS34-IN1 alone, *IFNB1* mRNA expression was increased 15-fold over control cells (Fig. 1E). This increase was accompanied by the phosphorylation of the transcription factor STAT1 at Tyr^701^ (P-STAT1) (Fig. 1F). P-STAT1 is a well-established downstream target of the type-I interferon receptor (IFNAR) signaling cascade and is essential for the expression and amplification of interferon-stimulated genes (ISGs) (*27*). As expected, CL307 stimulation induced robust levels of *IFNB1* mRNA transcription (a 60-fold increase). However, this response was attenuated in the presence of VPS34-IN1, along with a clear decrease in P-STAT1 levels (Fig. 1E and 1F). *IFNB1* mRNA transcription in response to CL307 was even more altered by spautin-1 (Fig. 1G), an alternative VPS34 inhibitor that promotes its degradation (*28*). These findings emphasize the indispensable role of VPS34 activity in regulating *IFNB1* expression in response to TLR7 agonists.

VPS34 inhibition had however a synergistic effect on the expression of the pro-inflammatory cytokine *TNF* mRNA (Fig. 1H), suggesting that alterations in PtdIns(3)P synthesis influence the balance between different TLR7 signaling pathways, favoring *TNF* mRNA expression (*18, 29*). When we monitored cytokine production by enzyme-linked immunosorbent assay (ELISA), we found that IFN-ß secretion was only detectable in response to CL307, while VPS34-IN1 treatment prevented it, leaving however TNF-α release unaffected (Fig. 1I). Thus, VPS34-IN1 alone triggers modest type-I IFN mRNA expression and STAT1 phosphorylation in CAL-1 at steady state, while shifting TLR7 signaling pathways towards a TNF-α rather than *IFNB1* expression mode after CL307 stimulation. Importantly, VPS34-IN1 appears to play a specific regulatory role in IFN synthesis or secretion, as the protein was poorly detected in the presence of the drug, regardless of its mRNA expression.

### VPS34-IN1 inhibits both the mTORC1/S6K and SGK3 signaling axis downstream of TLR7

We next monitored TLR7 expression and found that adding VPS34-IN1 to cells exposed to CL307 leads to a significant reduction in TLR7 levels (Fig. 2A). A reduction of type-I IFN production and STAT1 activation in treated cells (Fig. 1F), may reflect inhibition of a positive feedback loop that normally would drive TLR7 up-regulation, thereby preventing full cell activation and associated transcription.

**Figure 2.**
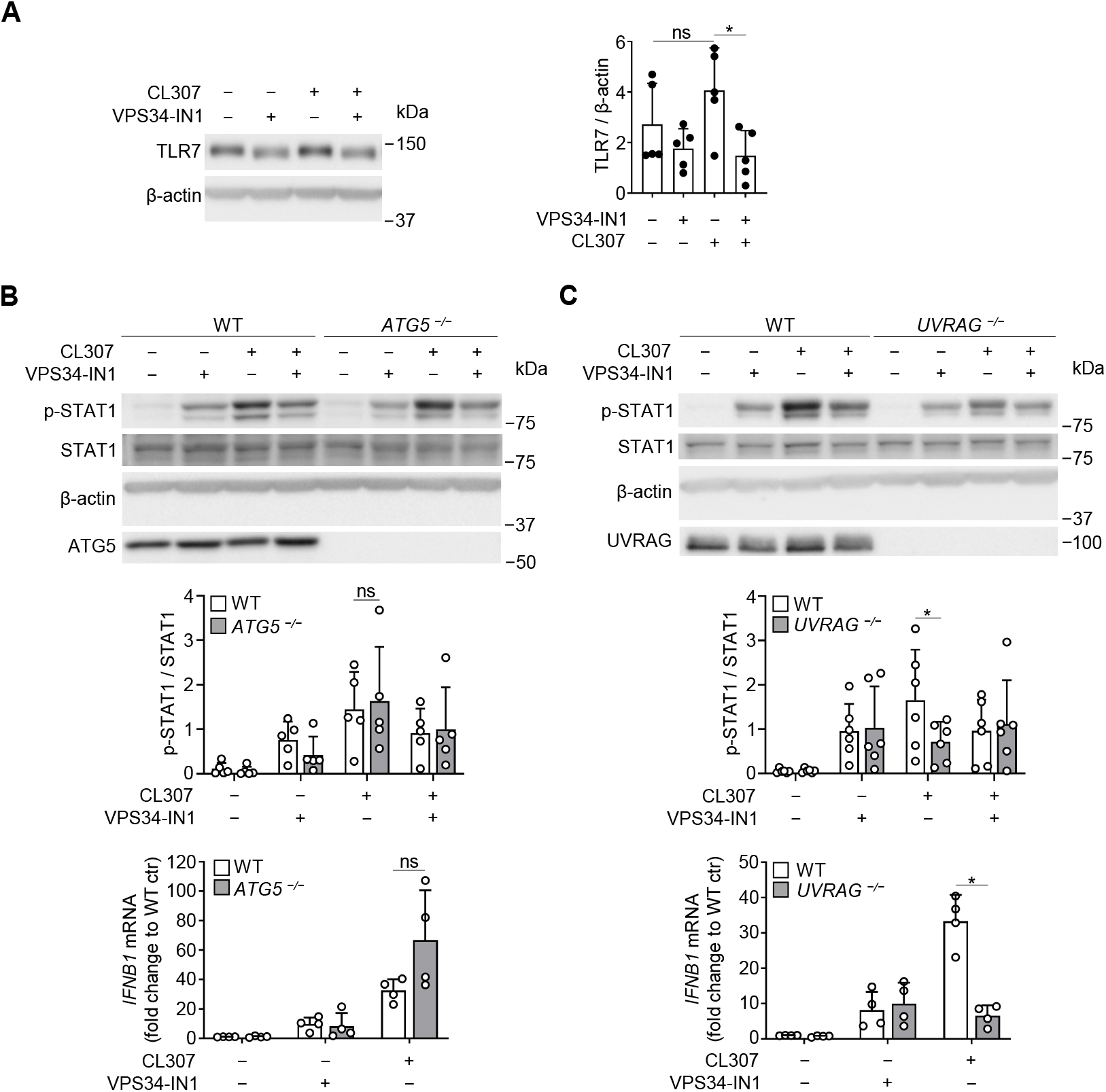
VPS34-IN1 affects TLR7 signaling by disrupting VPS34 complex II. (**A**) TLR7 was analyzed by immunoblot in CAL-1 cells subjected to CL307 and VPS34-IN1. (**B**, **C**) Levels of phosphorylated (Tyr^701^) STAT1, total STAT1, ATG5 and UVRAG were monitored for wild type (WT), *ATG5*^−/−^ and *UVRAG*^−/−^ CAL-1 cells treated with CL307 and/or VPS34-IN1. *IFNB1* mRNA levels were normalised to housekeeping gene (*GAPDH*) and compared to non-stimulated WT cells. (**A**-**C**) Bars show the mean ± SD and each dot shows data from one experiment. ns, non-significant results, *P < 0.05, by one-way ANOVA with Šidák post hoc test (A) and paired Student’s t test (B, C).

Recognizing the pivotal role of VPS34 in both autophagy and endosomal sorting (graphical abstract Fig. EV1A), we employed CRISPR/Cas9 technology to generate CAL-1 cells deficient in the *ATG5* and *UVRAG* genes (Fig. EV3A and EV3B). Subsequently, we characterized these selected mutants, confirming their expected deficits in LC3/ATG8 autophagy flux (*ATG5*^−/−^, Fig. EV3C) and the distribution of EEA1^+^ early endosomes (*UVRAG*^−/−^, Fig. EV3D). These mutant cells were then subjected to co-treatment with CL307 and VPS34-IN1 to assess the respective contributions of these pathways to TLR7 signaling. P-STAT1 levels and *IFNB1* mRNA levels were found similar among steady state and CL307 activated *ATG5*^−/−^ or control CAL-1 (Fig. 2B). A strong decrease in *IFNB1* mRNA induction, together with reduced STAT1 phosphorylation levels, were however observed in CL307-stimulated *UVRAG*^−/−^ cells (Fig. 2C). These results highlight that the disruption of the PI3KC3-CII complex through UVRAG deficiency and not of CI leads to a qualitative alteration in TLR7 signaling. This alteration involves the regulation of PtdIns(3)P production, which emerges as a critical factor controlling the expression of type-I IFN in response to TLR7 signaling in CAL-1.

TLR activation is well-documented to induce significant metabolic alterations upon dendritic cell (DC) activation, primarily through the stimulation of the AKT/mTORC1/S6K pathway, thereby augmenting various cellular responses, including enhanced protein synthesis (Fig. EV1B) (*30*,*31*). To evaluate the impact of these events, we employed flow cytometry to monitor protein synthesis using puromycylation detection (*32*). Strikingly, co-treatment with VPS34-IN1 resulted in the robust inhibition of mRNA translation (Fig. 3A). This inhibition suggests a disruption in the mTORC1/S6K signaling axis, as evidenced by the loss of phosphorylation in small ribosomal S6 and 4E-BP1 proteins upon VPS34-IN1 administration (Fig. 3B). Hence, VPS34-IN1 affects the capacity of TLR7 to activate the mTORC1/S6K pathway, partly through the inhibition of the PI3KC3-CII complex, as previously suggested in the case of amino acids and glucose starvation (*33*,*34*). This hypothesis gains further support from our observation of VPS34-IN1-induced inhibition of glycolysis and its subsequent enhancement upon TLR7 stimulation, as evidenced by a reduction in glucose consumption and the concomitant decrease in lactate production, quantified via NMR analysis in both CAL-1 cells and culture supernatants (fig. EV4A).

**Figure 3.**
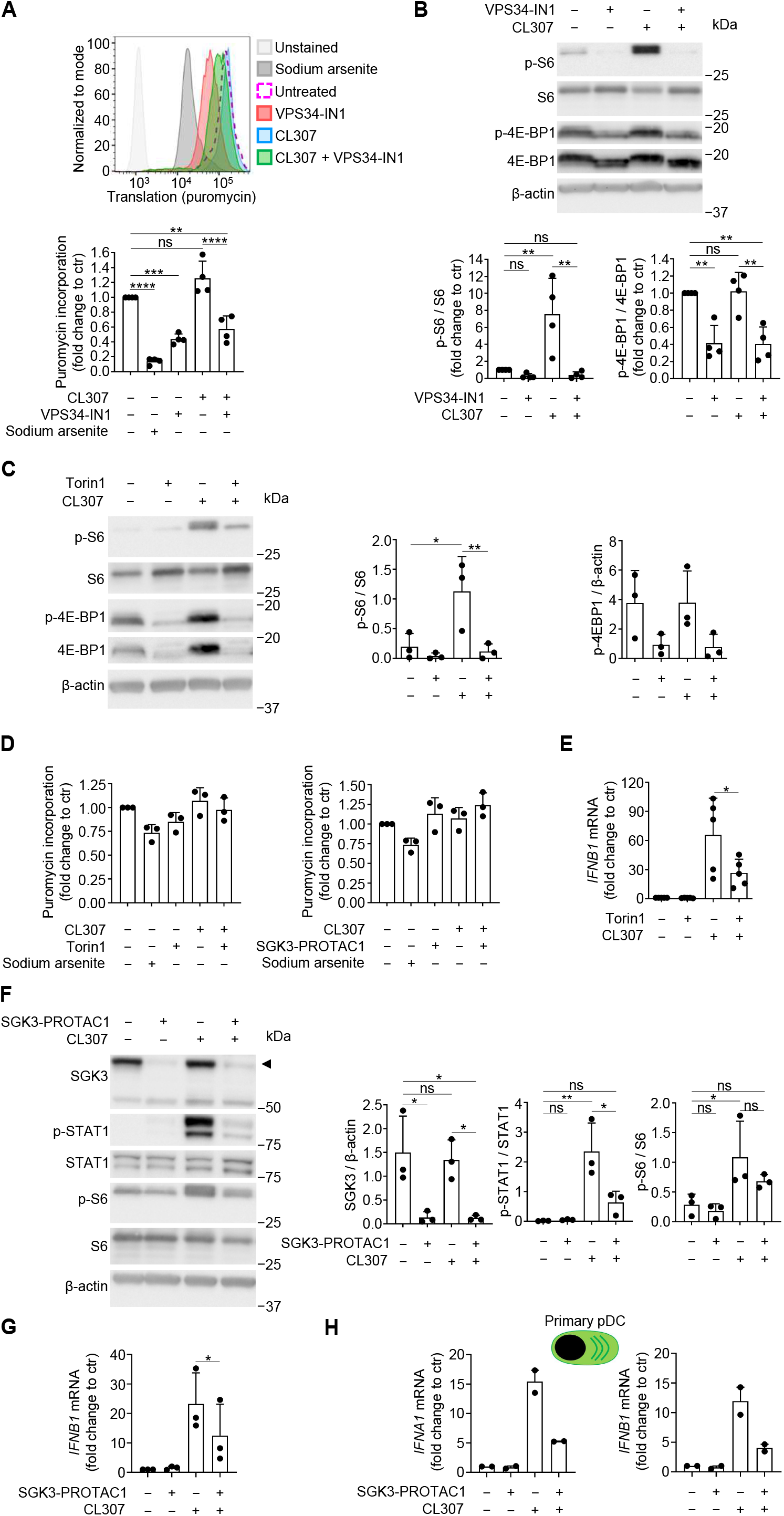
VPS34-IN1 inhibits both the mTORC1/S6k and SGK3 signaling axis downstream of TLR7. (**A,B**) CAL-1 cells were treated with CL307 and VPS34-IN1. (**A**) Protein synthesis was measured by flow cytometry upon puromycin incorporation. Sodium arsenite was used as negative control for puromycin incorporation. (**B**) Levels of total and phosphorylated S6 (Ser^235/236^) and 4E-BP1 (Ser^65^) were monitored by immunoblot. (**C**-**E**) CAL-1 cells were stimulated with CL307 and torin1 and levels of total and phosphorylated S6 (Ser^235/236^) and 4E-BP1 (Ser^65^) (**C**); protein synthesis (**D**) and *IFNB1* mRNA (**E**) were measured. (**F**, **G**) CL307-activated CAL-1 were treated with SGK3-PROTAC1 and levels of SGK3, p-STAT1 (Tyr^701^), STAT1, p-S6 (Ser^235/236^) and S6 and *IFNB1* mRNA expression were monitored. (**H**) Peripheral blood human pDCs were stimulated with CL307 and SGK3-PROTAC1 and levels of *IFNA1* (top) and *IFNB1* (bottom) mRNA were analyzed. (**A**-**H**) Bars show the mean ± SD and each dot shows data from one experiment. ns, non-significant results; *P < 0.05, **P < 0.01, ***P < 0.001, ****P < 0.0001, by one-way ANOVA with Tukey post hoc test (A, B, D, F), one-way ANOVA with Šidák post hoc test (C) and paired Student’s t test (E, G)).

mTORC inhibition was previously shown to suppress TLR3-induced IFN-ß production and mTORC1/mTORC2 to be direct targets of TBK1 for activation at Ser^2159^ (*35*). We reasoned that inhibition of the mTORC/AKT axis could be the main cause of protein synthesis alteration by VPS34-IN1, potentially affecting TLR7 and IFN-ß synthesis in activated pDC, as already suggested for TLR9 (*36*). CAL-1 activation was thus monitored upon treatment with 250 nM Torin1. The drug inhibited efficiently both S6 and 4E-BP1 phosphorylation (Fig. 3C), and slightly inhibited protein synthesis (Fig. 3D), while *IFNB1* expression (Fig. 3E) was dampened, as obtained with VPS34-IN1. The inhibitory effect of VPS34-IN1 on TLR7-mediated *IFNB1* expression appears therefore to be partially due to an effect on mTORC1 signaling by altering PtdIns(3)P synthesis and endo/lysosomes flux, preventing mTORC1 recruitment to and signaling from lysosomes (*37*).

Class I PI3Ks can activate isoforms of serum and glucocorticoid regulated kinases (SGKs), which share ∼50% identity within their catalytic domains to AKT and also control proliferation and survival responses in cancer cells (*38*). Among SGKs, SGK3, which possesses a PtdIns(3)P-binding PX domain, has been shown to substitute for AKT by phosphorylating TSC2 to activate mTORC1. The UVRAG/VPS34 complex can promote SGK3 recruitment and activation on endosomes, a process known to be inhibited by VPS34-IN1 treatment (*39*). We therefore used the SGK3-PROTAC1 degrader to silence specifically SGK3 in CAL-1 cells (*40*) and monitor the importance of this kinase for TLR7 signaling. Efficient SGK3 silencing induced a strong inhibition of STAT1 phosphorylation (Fig. 3F) and of type-I IFN mRNA expression (Fig. 3G) in response to CL307 stimulation. The phosphorylation of S6 that was observed with CL307 treatment was not as pronounced in the absence of SGK3 (Fig. 3F), even though there was no inhibition of protein synthesis (Fig. 3D). These results suggest that several redundant pathways regulating protein synthesis co-exist in CAL-1, a situation previously observed for AKT and SGK3 in the breast cancer lines ZR-75-1 and CAM-1 (*38*). The inhibitory effect of SGK3 inactivation on type-I IFN induction further suggests that in CAL1, SGK3 activity on molecules like STX7, STX12 or ITSN1, which all regulate endosomes dynamics (*41*), could be required for efficient TLR7 signaling.

We could further show that SGK3 inactivation also inhibited type-I IFN expression in response to CL307 in purified primary human blood pDCs (Fig. 3H). SGK3 is therefore a key actor of the downstream signaling of TLR7 and its inhibition by VPS34-IN1 likely contributes to decreased TLR7 signaling upon PtdIns(3)P synthesis inhibition (Fig. EV1E).

### STING is required for type-I IFN expression induction upon VPS34 inhibition

During our investigation, we observed that exposure to VPS34-IN1 alone had the ability to activate CAL-1 cells, as evidenced by IκBα degradation and concurrent expression of type-I IFN mRNA, independently of TLR7 stimulation (Fig. 4A). Several viral sensing networks, including the cGAS/STING and RLR/MAVS signaling pathways (Fig. EV1C), are known to trigger type-I IFN expression upon alteration of autophagy or membrane traffic (*42*,*43*). We therefore tested the impact of signaling adaptors MAVS or STING deletion (Fig. EV5A and EV5B) on the capacity of CAL-1 to express *IFNB1* in response to VPS34-IN1 treatment (Fig. 4B and 4C). As expected, MAVS deletion strongly impaired CAL-1 response to dsRNA (Poly(I:C), Fig. 4B). However, it had no effect on *IFNB1* expression nor STAT1 phosphorylation in response to VPS34-IN1 or CL307 (Fig. 4C). On the other hand, STING ablation completely inhibited the expression of *IFNB1* mRNA and subsequent STAT1 phosphorylation upon VPS34-IN1 treatment, while the response to CL307 remained unaffected (Fig 4D).

**Figure 4.**
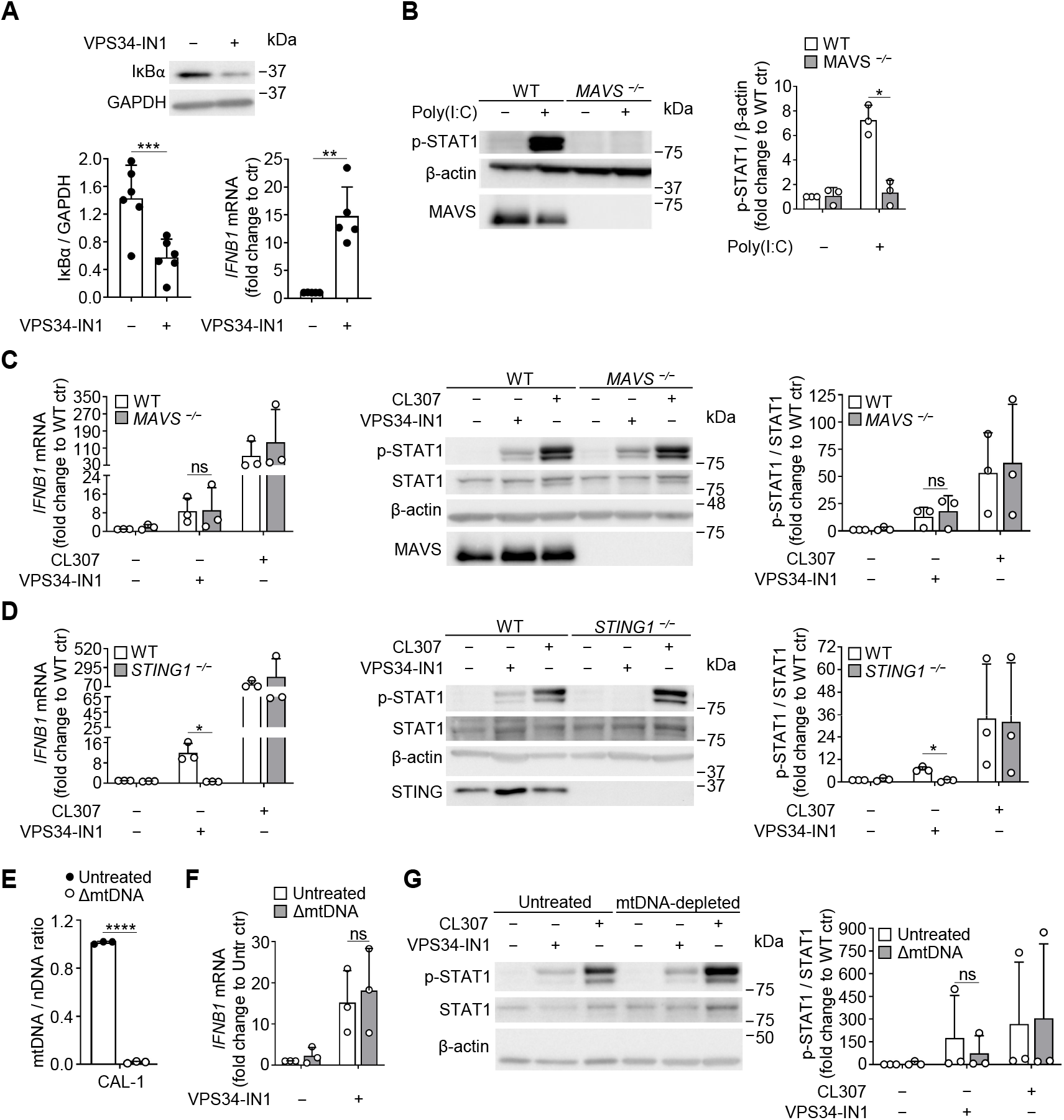
STING is required for type-I IFN expression induced by VPS34-IN1. (**A**) CAL-1 cells were treated with VPS34-IN1 and levels of IκBα and *IFNB1* mRNA were monitored by immunoblot and qPCR (**B**) Wild type (WT) and *MAVS*^−/−^ CAL-1 cells stimulated with Poly(I:C) were subjected to immunoblotting for phosphorylated STAT1 (Tyr^701^) and MAVS. (**C, D**) WT, *MAVS*^−/−^ and *STING*^−/−^ CAL-1 cells were treated with VPS34-IN1 or CL307 and *IFNB1* mRNA expression and levels of p-STAT1 (Tyr^701^), STAT1, MAVS and STING were monitored. (**E**) Levels of mitochondrial DNA (mtDNA) were monitored by quantifying levels of *MT-CO2* in relation to *GAPDH* in CAL-1 cells treated with 2’,3’-dideoxycytidine (ΔmtDNA). (**F**) Levels of *IFNB1* mRNA and (**G**) p-STAT1 (Tyr701) and STAT1 were monitored in cells treated with VPS34-IN1 or CL307, previously exposed to 2’,3’-dideoxycytidine (mtDNA-depleted) or untreated. (**A-G**) Bars show the mean ± SD and each dot shows data from one experiment. ns, non-significant results; *P < 0.05, **P < 0.01, ***P < 0.001, ****P < 0.0001, by paired Student’s t test.

Given VPS34-IN1 capacity to increase mitochondrial damage and ROS production (Fig. 1B), we wondered whether VPS34-IN1-induced release of mitochondrial DNA (mtDNA) could drive cGAS and STING activation (*44*). We efficiently depleted mitochondrial DNA using 2’,3’-dideoxycytidine (*45*) from CAL-1 cells exposed to VPS34-IN1 (Fig. 4E). Expression of *IFNB1* (Fig. 4F), and phosphorylated STAT1 (Fig. 4G) was found identical to control cells, ruling out an implication of abnormal mtDNA release in inducing cGAS/STING in VPS34-IN1-treated cells.

We next examined whether STING pre-activation by VPS34-IN1 could also potentiate its response to 2’3’-cGAMP (*46*). CAL-1 cells pre-treated with VPS34-IN1 for 1h, displayed hallmarks of a strong amplification of STING signaling with increased phosphorylation of STING (Ser^366^), TBK1 (Ser^172^) and IRF3 (Ser^396^) upon 2’3’-cGAMP delivery (*47*), which were all absent in *STING1*^−/−^ cells (Fig. 5A). The synergy between 2’3’-cGAMP and VPS34-IN1 resulted in a 40-fold enhancement of *IFNB1* mRNA expression (Fig. 5B) and an 8-fold increase in IFN-ß protein secretion (Fig. 5C). This synergistic effect was observed despite the reduction in protein synthesis caused by VPS34-IN1 (Fig. 5D) and dephosphorylation of S6 and 4E-BP1 (Fig. 5E).

**Figure 5.**
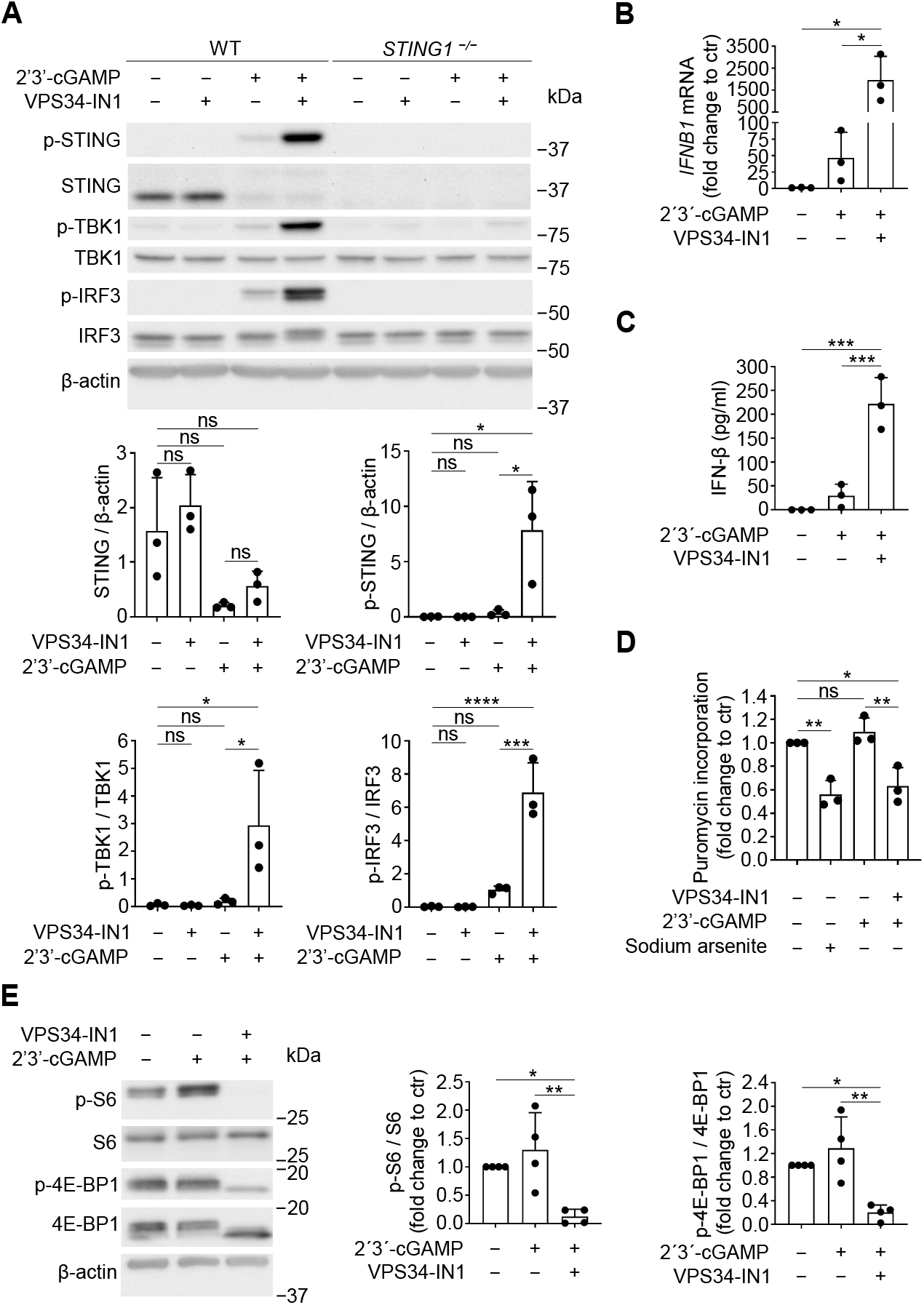
VPS34-IN1 intensifies STING-mediated IFN-β production triggered by 2’3’-cGAMP. (**A**) Wild type (WT) and *STING1*^−/−^ CAL-1 cells were treated with 2’3’-cGAMP and VPS34-IN1. Levels of total and phosphorylated forms of STING (Ser^366^), TBK1 (Ser^172^) and IRF3 (Ser^396^) were detected by immunoblot. (**B-E**) CAL-1 cells were treated with 2’3’-cGAMP and VPS34-IN1 and levels of *IFNB1* mRNA (**B**), secreted IFN-β (**C**), puromycin incorporation (**D**) as well as total and phosphorylated S6 (Ser^235/236^), and 4E-BP1 (Ser^65^) (**E**) were monitored. (**A-E**) Bars show the mean ± SD and each dot shows data from one experiment. ns, non-significant results; *P < 0.05, **P < 0.01, ***P < 0.001, ****P < 0.0001, by one-way ANOVA with Tukey post hoc test (A, D) and one-way ANOVA with Šidák post hoc test (B, C, E).

Pharmacological inhibition of VPS34 promotes STING activation and IFN-ß synthesis in synergy with 2’3’-cGAMP detection. STING activation was previously shown to also promote LC3 lipidation, in a ULK1- and VPS34-independent manner, driving conjugation of Atg8 proteins to Single Membranes (CASM) and remaining dependent on the ATG5/ATG16L1(WD40) lipid conjugation machinery (*48*). To clarify how VPS34-IN1 could amplify STING signaling, we monitored P-STING levels (Ser^366^) and *IFNB1* mRNA induction in CAL-1 cells inactivated for *ATG5 and UVRAG.* Upon 4h of 2’3’-cGAMP stimulation, STING degradation, known to be associated with its activation, together with TBK1 and IRF3 phosphorylation, was observed without significant differences between control, *ATG5*^−/−^ or *UVRAG*^−/−^ cells (Fig. 6A-B) and as anticipated on *IFNB1* induction by 2’3’-cGAMP (Fig. 6C-D). Co-treatment with VPS34-IN1 strikingly increased STING, TBK1 and IRF3 phosphorylation levels (Fig. 6A-B) in all cell types, as well as equivalently potentiated *IFNB1* expression (Fig. 6C-D). Thus, inhibition of macroautophagy, CASM and/or UVRAG dependent-processes (*44*,*49*,*50*), are not individually sufficient to trigger the potentiating effect of VPS34-IN1 on 2’3’-cGAMP-dependent signaling in CAL-1 cells.

**Figure 6.**
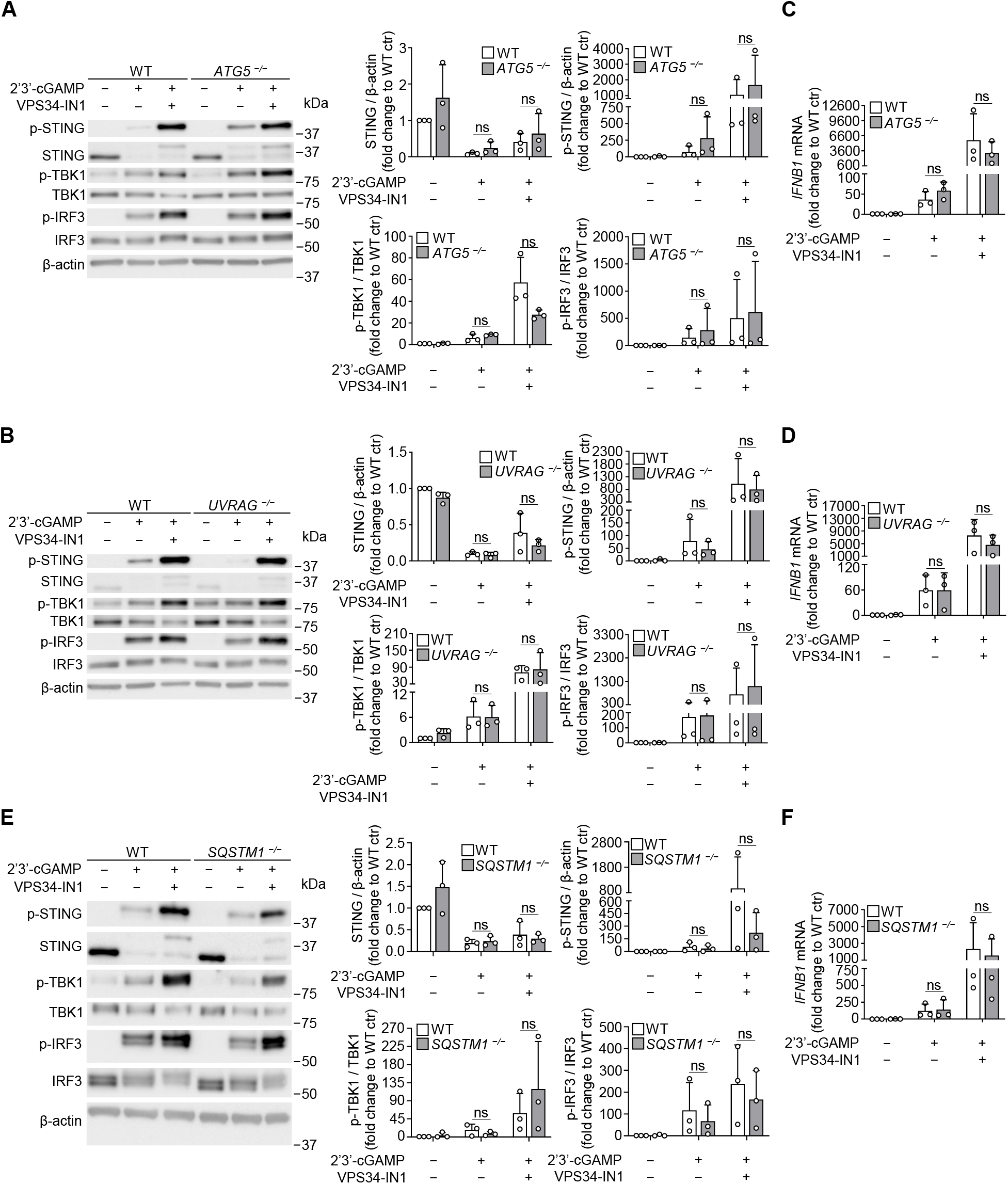
Potentiation of STING signaling by VPS34-IN1 does not require macroautophagy or UVRAG-dependent processes. (**A-F**) Wild type (WT), *ATG5*^−/−^, *UVRAG*^−/−^ and *SQSTM1*^−/−^ CAL-1 cells were stimulated with 2’3’-cGAMP and VPS34-IN1. Total and phosphorylated levels of STING (Ser^366^), TBK1 (Ser^172^) and IRF3 (Ser^396^) were analyzed by immunoblot (**A, B, E**) and *IFNB1* mRNA levels by qPCR (**C, D, F**). (**A**-**F**) Bars show the mean ± SD and each dot shows data from one experiment. ns, non-significant results by paired Student’s t test.

TBK1-dependent phosphorylation of the selective autophagy receptor p62/SQSTM1 was shown to be essential for cGAMP-stimulated degradation of STING in mouse embryonic fibroblasts (MEF) and THP-1 cells (*51*). Consequently, STING was described not to be degraded in *p62*/*SQSTM1*-deficient cells, which in turn produce augmented levels of type-I IFN. We tested whether p62 inactivation (Fig. EV6A) could have more severe consequences than that of *ATG5* in CAL-1 cells. Similar to control cells, 2’3’-cGAMP stimulation of *p62/SQSTM1*^−/−^ cells induced a major loss of STING, a modest accumulation of p-STING and a strong synergy with VPS34-IN1, leading to an equivalent enhancement of TBK1 and IRF3 activation (Fig. 6E). These observations were confirmed by matched levels of *IFNB1* induction in the different cell lines tested upon 2’3’-cGAMP delivery in synergy or not with VPS34-IN1 (Fig. 6F). We also investigated the role of the ULK1 kinase, which is itself inhibited by mTORC1-mediated phosphorylation (*52*) and, in addition to its key role in regulating autophagy, has been proposed to control STING activity through direct phosphorylation at Ser^366^ (*42*). ULK1 inactivation inhibited the LC3 autophagy flux in normal and nutrient starvation conditions (EBSS, fig S6B and S6C). Despite this deficiency, no difference in the capacity of *ULK1*^−/−^ and control CAL-1 to amplify STING signaling and I*FNB1* expression in response to 2’3’-cGAMP and VPS34-IN1 co-treatment was observed (fig S6D and S6E). Thus, differently from other cell types (*51*), STING signaling potentiation by *p62*, *ATG5* or *ULK1* deletion was not observed in CAL-1, confirming that autophagy inhibition is not the prime cause of activated STING stabilization and enhanced type-I IFN production observed after VPS34-IN1 treatment.

### VPS34-IN1 drives cap-mediated translation inhibition and PPP1R15A/GADD34 synthesis independently of the ISR

Importantly, cell-autonomous responses involving PERK triggering by STING have been shown to regulate translation of viral and host proteins (*53*), as well as to protect cell integrity from the co-lateral damages linked to STING activation (*54,55*,*56*). In regards to the protein synthesis inhibition observed upon VPS34-IN1 treatment, we investigated the capacity of the drug to trigger an ER stress unfolded protein response (UPR) by altering the lipid composition and dynamics of ER membranes targeted by the PI3KC3-CI complex (*57*) (Fig. EV1D). ER stress could potentially decrease protein synthesis through the induction of the integrated stress response (ISR) or the IRE1-dependent branch of the UPR (*58*), as well as promote type-I IFN and cytokines expression (*59*). Activation of the ER-resident eIF2AK3 (PERK), and phosphorylation of its major target, the eukaryotic initiation factor 2-α (eIF2-α), were not observed during a time dependent response to VPS34-IN1 (Fig. 7A), confirming that ISR induction is not responsible for translation repression. Additionally, IRE1α-dependent splicing of the key UPR transcription factor XBP1 mRNA was not induced by VPS34-IN1 treatment (Fig. 7B), confirming that it does not activate two of the main UPR signaling branches and is therefore unlikely to promote important ER stress and associated loss of protein synthesis in CAL-1 cells.

**Figure 7.**
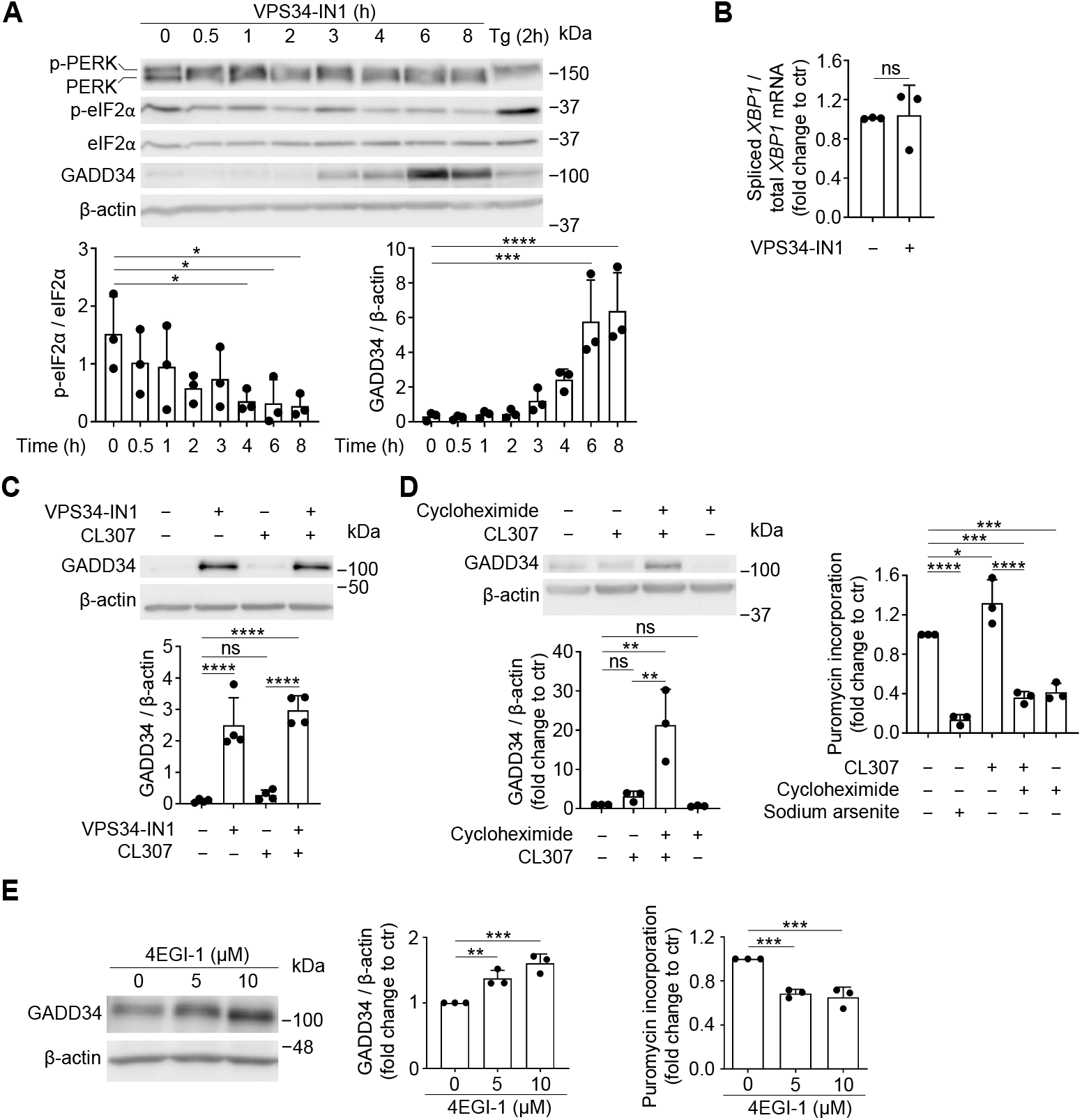
VPS34-IN1 promotes GADD34 synthesis independently of the ISR. (**A**) Levels of PERK, total and phosphorylated (Ser^51^) eIF2α and GADD34 were monitored for CAL-1 cells treated for different time points with VPS34-IN1. Thapsigargin (Tg) treatment was used as positive control. (**B**) Spliced *XBP1* mRNA levels were normalized to total *XBP1* mRNA. (**C**) CAL-1 cells were treated with CL307 and VPS34-IN1 (8 h) and GADD34 production was monitored by immunoblot. (**D**) Cells were activated with CL307 and cycloheximide and levels of GADD34 and puromycin incorporation were detected by immunoblot or flow cytometry, respectively. (**E**) CAL-1 cells were treated with 4EGI-1 and levels of GADD34 and puromycin incorporation were monitored. (**A-E**) Bars show the mean ± SD and each dot shows data from one experiment. ns, non-significant results; *P < 0.05, **P < 0.01, ***P < 0.001, ****P < 0.0001, by one-way ANOVA with Dunnett post hoc test (A, E), paired Student’s t test (B) or one-way ANOVA with Tukey post hoc test (C, D).

Although increased eIF2-α phosphorylation and ATF4 synthesis are normally a requisite for the synthesis of the PP1c phosphatase adaptor PPP1R15a (GADD34) (*60*), a strong accumulation of this negative feedback regulator of the ISR (*61*) was observed in VPS34-IN1 treated cells (Fig. 7A and 7C). GADD34 is known to positively impact *IFNB1* expression in virally infected MEFs and activated mouse DCs (*62*); (*63*). GADD34 synthesis upon VPS34-IN1 treatment was found independent of an observable increase in eIF2-α phosphorylation, differently from a *bona fide* ISR induction by thapsigargin (Tg) (Fig. 7A). GADD34 synthesis although translationally regulated by small decoy upstream open reading frames (uORFs) (*64*), can occur independently of eIF2-α phosphorylation in situations during which translation intensity is tuned-down or qualitatively affected, as seen with VPS34 inhibition. Cycloheximide (CHX) was used to inhibit protein synthesis in CAL-1 cells and further demonstrates that GADD34 accumulation is indeed correlated with the magnitude of translation inhibition when cells are activated with CL307 (Fig. 7D). VPS34-IN1 treatment likely results in inhibition of cap-dependent translation by increasing 4E-BP1 high affinity binding to eIF4E and interfering with eIF4G and ultimately eIF4F complex recruitment to m7G capped-mRNAs. We therefore tested if pharmacological inhibition with 4EGI-1 (*65*) could recapitulate the consequences of VPS34-IN1 treatment during CAL-1 activation. Upon 5h exposure to 4EGI-1, expression level of GADD34 increased while leading to protein synthesis inhibition (Fig. 7E), confirming that eIF4G1 inhibition could mimic the impact of VPS34-IN1 on mRNA translation both quantitatively and qualitatively.

### Cap-dependent translation inhibition triggers STING signaling potentiation by VPS34-IN1

Our results support that alteration in transport or degradation alone or protein synthesis inhibition alone are insufficient to drive STING signaling upon VPS34-IN1 exposure. Transient cap-mediated translation inhibition is however likely to potentiate STING activation and type-I IFN expression, through the down-regulation of neosynthesized negative feedback signaling regulators of anti-viral pathways (*62*). We therefore tested if the VPS34-IN1-mediated translation inhibition could potentiate STING signaling using CHX or 4EGI-1. Amplification of STING signaling was recapitulated after co-treatment with 2’3’-cGAMP with both inhibitors (Fig. 8A and 8B). Importantly, *IFNB1* mRNA levels were strongly enhanced by both co-treatments, mirroring VPS34-IN1 effect (Fig. 8C). Translation alteration by VPS34-IN1 via inhibition of 4E-BP1 phosphorylation is therefore likely to be the major cause for the strong type-I IFN induction observed upon co-treatment with 2’3’-cGAMP. Differently from TLR7 stimulation, SGK3 silencing in cells co-stimulated or not with 2’3’-cGAMP had no impact on STING signaling nor *IFNB1* mRNA expression, as anticipated from its lack of effect on protein synthesis in CAL-1 cells (Fig. 8D).

**Figure 8.**
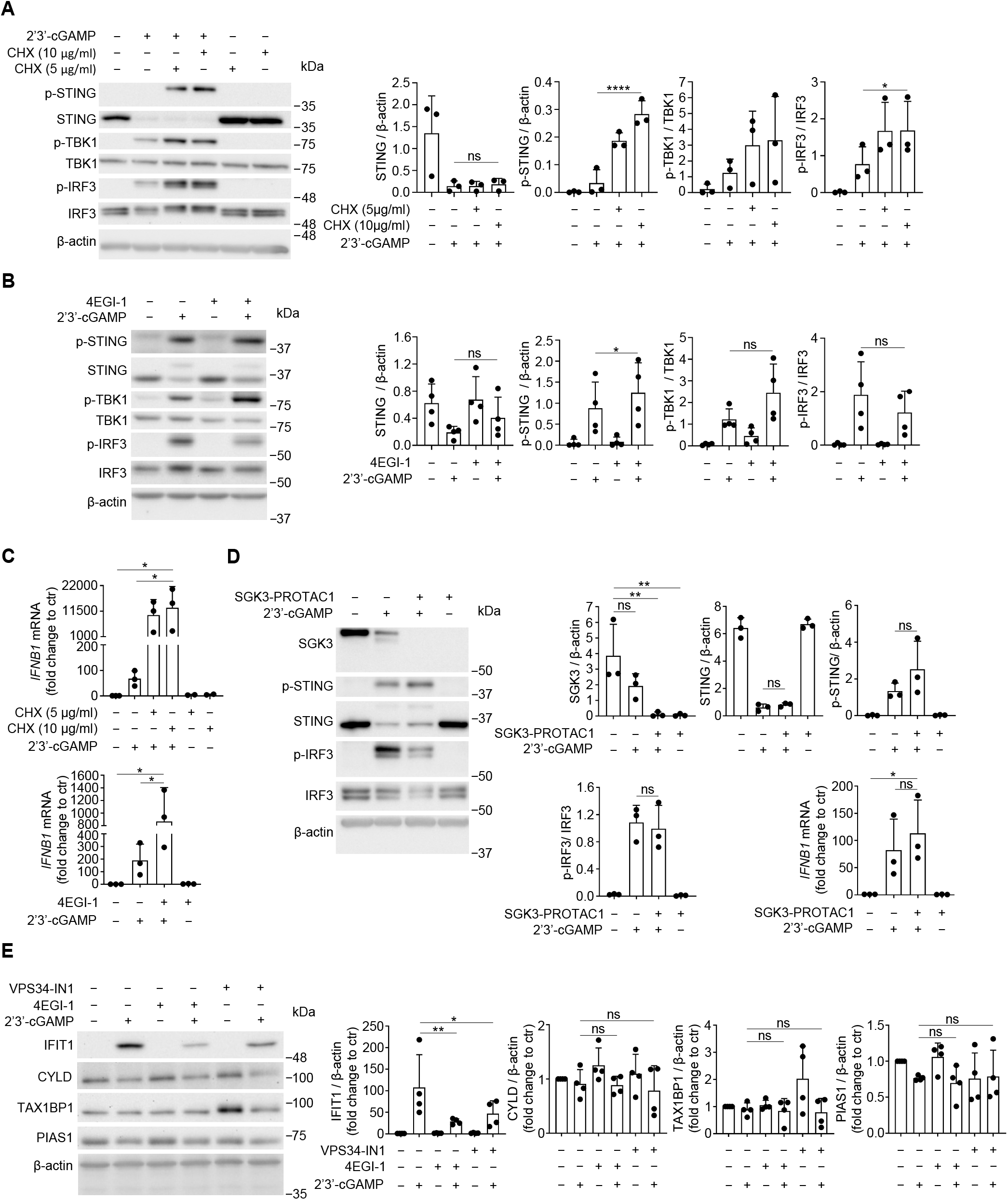
Inhibiting cap-dependent translation boosts STING signaling. (**A-D**) CAL-1 cells were stimulated with 2’3’-cGAMP, cycloheximide (CHX), 4EGI-1 and SGK3-PROTAC1. Levels of total and phosphorylated STING, TBK1 and IRF3 (Ser^366^, Ser^172^ and Ser^396^, respectively), of total SGK3 (D), as well as *IFNB*1 mRNA (C, D) were monitored. (**A, C, D**) Cells were pre-treated for 1h with either CHX, 4EGI-1 or SGK3-PROTAC1, followed by 4h incubation with 2’3’-cGAMP. (**B**) Cells were pre-treated for 1h with 4EGI-1 followed by 1h incubation with 2’3’-cGAMP. (**E**) Cells were treated with 2’3’-cGAMP, 4EGI-1 and VPS34-IN1 and levels of IFIT1, CYLD, TAX1BP1 and PIAS1 were monitored by immunoblot. (**A**-**E**) Bars show the mean ± SD and each dot shows data from one experiment. ns, non-significant results; *P < 0.05, **P < 0.01, ****P < 0.0001, by paired Student’s t test (A, B), by one-way ANOVA with Šidák post hoc test (C, E), by one-way ANOVA with Dunnett post hoc test (SGK3) or paired Student’s t test (remaining proteins) and by one-way ANOVA with Šidák post hoc test (D).

As anticipated from our previous work on MEFs (*66*), the levels of several negative feed-back regulators of innate and cytokine receptors transduction pathways were examined upon VPS34-IN1 treatment. Strikingly, the levels of IFIT1/ISG56 were strongly inhibited by VPS34-IN1 or 4EGI-1 during cGAMP stimulation (Fig. 8E). Interestingly, ISG56 has been shown to bind STING and disrupt its interaction with TBK1, leading to the inhibition of virus-induced IRF3 activation, *IFNB1* expression, and cellular antiviral responses (*67*). Thus, by triggering STING activation and inhibiting cap-mediated translation of negative feedback regulators such as ISG56, VPS34-IN1 strongly enhances STING signaling and *IFNB1* expression (*66*).

Overall, the effects of VPS34-IN1 on TLR7 and on STING signaling are triggered by distinct biochemical pathways, SGK3 inhibition being dominant for the former, while cap-mediated translation inhibition being key for STING signaling potentiation.

## Discussion

VPS34, as the catalytic subunit of the phosphatidylinositol 3-kinase (PI3K) class III complex, plays a major role in generating PtdIns(3)P that is necessary for endosomes trafficking and dynamics. VPS34 implication in many cellular functions linked to endocytosis and autophagy has led to the development of highly specific pharmacological inhibitors (*68*). These inhibitors, like VPS34-IN1, have been recently proposed as chemotherapeutic agents to treat leukemia or breast cancer (*11*,*12*). Indeed, systemic treatment of tumor-bearing mice with VPS34-IN1 induces a potent inflammatory signature in the tumor microenvironment, including activation of the STAT1/IRF7 pathway that is responsible for this anti-tumoral effect (*21*). VPS34 is also required for the maintenance of primary mouse naïve T cells, since its deletion causes an increase in damaged mitochondria mass and accumulation of reactive oxygen species, likely due to an impairment of autophagy in *VPS34* ^-/-^ T cells (*69*). Using an *in vitro* approach, we provide evidence that although VPS34-IN1 has a weak toxicity on leukemic blastic plasmacytoid dendritic cell neoplasm, it activates STING and can strongly synergize with 2’3’-cGAMP to produce high amounts of type-I IFN, resulting in STAT1 activation.

Inactivation of key genes in CAL-1, including *UVRAG*, *ATG5 and p62* has determined that inhibition of macroautophagy, CASM and/or UVRAG dependent-processes are not individually sufficient to prevent efficiently STING degradation and potentiate its 2’3’-cGAMP-dependent signaling in CAL-1 cells, differently from other cell types (*44,50*,*51*). This finding also contrasts with the recent implication of microautophagy and of VPS molecules in allowing STING degradation in MVBs (*70*). In MEFs, the knockdown of the PtdIns(3)P effector Tsg101, VPS34 or VPS15 is sufficient to prevent lysosomal degradation, termination of STING signaling, and exacerbated type-I IFN expression, a situation we did not observe upon *UVRAG* inactivation or VPS34 inhibition in CAL-1 cells. The STING ER exit protein (STEEP; also known as CxORF56) has been proposed to associate with STING after its activation, promoting COPII-dependent STING trafficking from ER by stimulating production of PtdIns(3)P and formation of ER membrane curvature (*71*). Remarkably, STEEP deficiency retains STING at ER and reduces STING-induced immune response in virus-infected cells and in cells from patients with STING-associated diseases (*71*). Likewise, sorting nexin 8 (SNX8) was reported to be decisive for STING transit from ER through recruitment of VPS34 (*43*), that is also recruited by STEEP. Moreover, PtdIns(3)P phosphatases myotubularin-related protein (MTMR)3 and MTMR4 suppress STING-mediated immune responses by influencing STING trafficking (*72*). All these pathways could therefore be affected by VPS34 inhibition and allow some level of STING activation. We cannot formally exclude that increased persistence of STING upon VPS34 inhibition does not contribute to enhanced signaling, but concomitant inhibition of cap-mediated translation potentiates STING signaling in CAL-1 cells. This also suggests that strong negative feed-back mechanisms controlling the different signaling pathways leading to type-I IFN production might pre-exist in non-activated pDCs.

VPS34-IN1 is known to inhibit the mTORC1 pathway in leukemic cells and during nutrient scarcity, as it regulates amino acid and glucose inputs to mTOR and S6K1 (*11*,*33*). SGK3, which shares roughly 50% of its catalytic domain identity with AKT (*73*), can also activate mTORC1, albeit partially in CAL-1 cells, possibly in conjunction with AKT or another SGKs. Notably, SGK3 has a selective PtdIns(3)P-binding PX domain that is highly sensitive to VPS34-IN1 and becomes inactive when PtdIns(3)P synthesis is disrupted (*8*). In pDCs, SGK3 participates to the signaling events leading to type-I IFN induction upon TLR7 stimulation, while not being involved in STING activation by cGAMP. This contrasts with previous observations with macrophages, where SGK3 inhibition diminishes IFN expression upon DMXAA stimulation (*74*). VPS34-IN1 also inhibited mTORC1 signaling, presumably by altering its recruitment to endo-lysosomes, and impacting on TLR7 signaling. The cumulated effects of VPS34-IN1 on these different pathways result in a strong loss of cap-mediated protein synthesis. Intriguingly, endosomal proteins like STX7, STX12, and ITSN1 are known SGK3 targets, and their phosphorylation enhances their ability to regulate endosome dynamics. VPS34 together with SGK3 activation likely controls endosome dynamics, creating favorable conditions for the assembly of the TLR7 signalosome and driving type-I IFN expression, rather than TNF-α in response to CL307.

VPS34-IN1 inhibits the phosphorylation of 4E-BP1 and promotes the translation of molecules like GADD34/PPP1R15A, a co-factor for PP1c typically expressed during the ISR. Dephosphorylation of 4E-BP1 has recently been shown to disrupt eIF1 interaction with eIF4G1, a protein complex promoting ribosome scanning and stringent AUG selection in translated mRNAs. This dissociation enhances the translation of ISR genes regulated by upstream open reading frames (uORFs), such as GADD34, independently of eIF2-α phosphorylation (*64*,*75*).

We have shown the treatment with the inhibitor eIF4GI-1 alone is sufficient to induce GADD34 synthesis, confirming that altering cap binding favors translation of uORF-regulated mRNAs. GADD34 regulates the expression of several cytokines, including IFN-ß and interleukin-6 (IL-6), in activated DCs (*76*). Therefore, it could also positively influence *IFNB1* expression upon exposure to VPS34-IN1. Importantly, the neo-synthesis of GADD34 indicates that VPS34-IN1 induces profound changes in the intensity and quality of translation mediated by activated CAL-1 cells. As demonstrated with CHX and 4EGI-1, reducing cap-mediated translation enhances GADD34 synthesis but can inhibit or delay the synthesis of key negative feedback regulators of TBK1, NF-kB, or STAT1 signaling, which in turn potentiates STING signaling and facilitates type-I IFN expression. Negative feedback regulators generally depend on neosynthesis for their expression. The amplification of antiviral pathways and *IFNB1* mRNA expression during the ISR has been previously demonstrated for RLRs and cytosolic dsRNA detection by PKR (*66*). This principle extends to other innate pathways requiring TBK1 activation. For example, pre-stimulation of STING in human pDCs induces the expression of SOCS1 and SOCS3, creating an autoinhibitory loop that impedes IFN production upon successive TLR9-mediated DNA recognition (*77*). Here, this situation is likely prevented by VPS34-IN1-mediated translation inhibition, which by targeting molecules like IFIT1/ISG56, amplifies the STING response in activated CAL-1 cells. The synthesis of negative feedback regulators in human pDCs is a part of the cross-talk between innate signaling pathways, essential for dampening IFN production and preventing accidental and uncontrolled cytokine release.

Taken together our observations suggest therefore that VPS34-IN1 induces significative STING and STAT1 activation in CAL-1 cells via protein synthesis inhibition which dominates over changes in PtdIns(3)P levels that have been shown to be generally inhibitory for STING activity. Other compounds capable of inhibiting cap-mediated translation are therefore likely to have a similar effect during chemotherapy or immunotherapy. While these compounds may lack strong cytotoxic effects on cancer cells, they could potentially exhibit anti-tumoral activity by promoting type-I IFN or pro-inflammatory cytokine production and transforming the tumor microenvironment, as recently suggested for VPS34-IN1 (*21*). However, VPS34-IN1’s impact on various cellular functions may present challenges for its clinical use. Additionally, our discovery that SGK3 is crucial for TLR7 signaling in pDCs opens new possibilities for developing SGK3 inhibitors to control autoimmune syndromes, often driven by excessive NA sensing and type-I IFN production (*78*).

## Methods

### Cells and reagents

CAL-1 human pDCs (gift from Dr. Takahiro Maeda, Nagasaki University, Japan) were grown in RPMI 1640 medium with 2 mM L-glutamine (Gibco), supplemented with 10% heat-inactivated fetal bovine serum (FBS) (Sigma-Aldrich), 1 mM sodium pyruvate, 10 mM HEPES, 1x non-essential amino acids (all from Gibco). Cells were maintained at 37°C and 5% CO_2_ and were routinely tested for mycoplasma contamination. In each experiment, cells were cultured overnight at 1 × 10^6^ cells/ml in a 24-well plate, 1 ml per well, in complete medium with 1% FBS. For starvation, EBSS with calcium and magnesium (Gibco) was used as medium.

Primary pDCs were purified from mononuclear cells enriched from healthy blood donors’ buffy coats (Établissement Français du Sang (EFS), Marseille, FR). Cells were submitted to density gradient centrifugation using Ficoll-paque plus (Cytiva, Marlborough, MA, USA) and pDCs were purified using a negative selection MACS beads-based assay (Plasmacytoid Dendritic Cell Isolation Kit II human, Miltenyi Biotec, Bergisch Gladbach, GER), following manufacturer’s instructions. The isolated pDCs (85% purity) were maintained overnight in the presence of IL-3 (1 ng/ml) (PeproTech, London, UK) before stimulation.

VPS34-IN1 (at 5 µM unless otherwise indicated, for a total incubation time of either 8h, when in combination with CL307 or 5h, when in combination with 2’3’-cGAMP) and 4EGI-1 (10μM, with a pre-incubation of 1h prior to cell activation for either 1 or 4h) were both from Cayman Chemical. CL307 (1 µM, 6 h), 2’3’-cGAMP (5 µg/ml, 4h) and Poly(I:C) (50 µg/ml, 6h) were from InvivoGen. Spautin-1 (Calbiochem) was used at 10 µM for 19h. SGK3-PROTAC1 (MedChemExpress) was used at 2 µM with a total incubation time of either 8h, when in combination with CL307, or 5h, when in combination with 2’3’-cGAMP. Cycloheximide (5 or 10 µg/ml, for a total incubation time of 8h when in combination with CL307, or 5h when in combination with 2’3’-cGAMP) was from Sigma-Aldrich. Bafilomycin (100 nM, 4 h) was from Merck Millipore. Thapsigargin (Santa Cruz Biotechnology) was used at 500 nM for 2h, and Torin1 (Bio-Techne) was used at 250 nM for 8h total incubation time, with 2h pre-incubation followed by 6h in combination with CL307. All used reagents are listed in Appendix Table S1.

### CRISPR/Cas9 knockout

Gene inactivation was performed with CRISPR/Cas9 technology in an inducible Cas9 expressing CAL-1 cell line, which constitutively expressed GFP. Guide RNAs (gRNA) were generated using the GeneArt Precision gRNA synthesis kit (Invitrogen). The genomic target sequences were searched by accessing the GeneArt CRISPR Search and Design Tool (thermofisher.com/crisprdesign) or were found in literature: 5’-AACTTGTTTCACGCTATATC-3’ for *ATG5* gene; 5’-ACGTGGCGAAGTCTCGATTT-3’ for *UVRAG*; 5’-GCTGGGACTGCTGTTAAACG-3’ for *STING1*; 5’-TCTGGGGCTGAGCGTCTGCA-3’ for *MAVS*; 5’-AAGTTCGAGTTCTCCCGCA-3’ for the *ULK1* gene; 5’-ATGGCCATGTCCTACGTGA-3’ for *SQSTM1*; 5’-GAGCTGGACGGCGACGTAAA-3’ for GFP. All gRNA target sequences are listed in Appendix Table S2. Cells to be gene-modified were cultured in the presence of 2 μg/ml doxycycline (Sigma-Aldrich) for 48 h to induce Cas9 production. For gRNA delivery, cells were washed and resuspended in 100 μl of RPMI 1640 medium at room temperature and then electroporated (875 V/cm, 950 μF and 25 Ω) in a 4 mm cuvette, using an ECM 630 electroporation system (BTX). The electroporation mixture included 6.25 × 10^5^ cells and 5 μg of each of two different gRNAs: one GFP-specific and another targeting the gene of interest. Cells were subsequently subjected to limiting dilutions and seeded on 96-well plates to obtain clonal cell populations. Single clones were analyzed by flow cytometry and the GFP-negative ones were selected for expansion and tested by immunoblotting and DNA sequencing (GATC Biotech, Germany) to confirm gene deletion efficiency.

### 2’3’-cGAMP transfection

For cGAMP treatment, 10^6^ cells were resuspended in 160 μl of RPMI 1640 medium and 2’3’-cGAMP was transfected at 5 µg/ml by electroporation (875 V/cm, 950 μF and 25 Ω) in a 4 mm cuvette, using an ECM 630 electroporation system (BTX). Finally, cells were transferred to a new 24-well plate with new supplemented medium.

### Depletion of mitochondrial DNA

To eliminate mtDNA, CAL-1 cells were treated for 5 days with 100 μM 2’,3’-dideoxycytidine (Alfa Aesar) in medium supplemented with 50 μg/ml uridine (Sigma-Aldrich). To confirm mtDNA depletion, total DNA was extracted using NZY Tissue gDNA Isolation kit (NZYTech). Three different DNA quantities (15, 7.5 and 3.75 ng) were used as template of Real-Time PCR (qPCR) for mitochondrial *MT-CO2* gene (forward primer 5’-CGTCTGAACTATCCTGCCCG-3’; reverse primer 5’-TGGTAAGGGAGGGATCGTTG-3’) and for the nuclear *GAPDH* gene (forward primer 5’-ATGCTGCATTCGCCCTCTTA-3’; reverse primer 5’-GCGCCCAATACGACCAAATC-3’). 2xSYBR Green qPCR Master Mix (Low ROX) (Bimake) was used to perform qPCR. Ratios of 2^−CT^ for mitochondrial MT-CO2 over nuclear GAPDH were averaged and the fold change relative to untreated cells was determined, as described by others (*45*).

### RNA extraction and qPCR

Total RNA was isolated using the RNeasy Mini Kit (Qiagen), including the DNase (RNase-Free DNase set, Qiagen) treatment to remove residual DNA. Superscript II reverse transcriptase (Invitrogen) was combined with 500 ng of total RNA, random hexamer primer (Thermo Scientific), dNTPs mix (NZYTech) and Ribonuclease inhibitor (NZYTech) for cDNA synthesis. qPCR reactions included a 2x SYBR Green qPCR Master Mix (Low ROX) (Bimake), 300 nM of each specific primer and 2 μL of diluted cDNA as template. The assays were performed on a 7500 Real-Time PCR system (Applied Biosystems). The qPCR conditions were the following: initial denaturation at 95 °C for 5 min; 40 cycles of denaturation at 95 °C for 15 s, as well as annealing and extension at 60°C for 60 s. To confirm the specificity of the assays, a melting curve was generated for every run. Gene expression was determined after normalization to *GAPDH* expression (housekeeping gene). Data are presented as fold change relative to control or to untreated WT cells, as indicated. All used primers are listed in Appendix Table S3.

### Protein extraction and immunoblotting

Proteins were extracted in Tris buffer (Tris 20 mM pH 8, NaCl 10 mM, MgCl_2_ 1.5 mM) 1% Triton X-100 (Sigma-Aldrich), supplemented with 1 tablet of protease inhibitors (Thermo Scientific), MG-132 (5 µM, CSNpharm) and phosphatase inhibitors (50 mM sodium fluoride and 0.2 mM sodium orthovanadate, both from Sigma). Protein concentration was measured using the BCA protein assay kit (Thermo Scientific). 8 to 25 μg of protein were separated by SDS-PAGE on 10% or 4-15% polyacrylamide gradient gels, which was followed by transfer onto PVDF membranes (Millipore) and blocking for 1 h at room temperature (RT), in Tris-buffer with 0.05% Tween 20 (Sigma) (TBST) (Sigma) and 5% bovine serum albumin (BSA) (NZYTech). Membranes were incubated overnight at 4°C with primary antibodies diluted in 1% BSA/TBST and were probed with secondary antibody for 1h at RT. Antibodies were detected in a ChemiDoc Imaging System (Bio-Rad) using ECL Select Western Blotting Detection Reagent (Cytiva), ECL Plus Western Blotting Substrate or ECL Western Blotting Substrate (both from Thermo Scientific) according to the protein abundance. Image Lab Software (Bio-Rad) was used for quantification of protein bands. Antibodies used in Western blot were (listed in Appendix Table S4): mouse monoclonal antibodies anti-β-actin and anti-UVRAG (both from Sigma-Aldrich), anti-LC3B (NanoTools), anti-IκB-α (Carboxy-terminal) (Cell Signaling Technology), anti-p-STAT1 (Tyr^701^) (BioLegend), anti-STAT1 (one from BioLegend, another from Santa Cruz Biotechnology), anti-p-4E-BP1 (Ser^65^) and anti-MAVS (all the previous from Santa Cruz Biotechnology); rabbit monoclonal antibodies anti-ATG5, anti-p-eIF2α (Ser^51^), anti-p-TBK1 (Ser^172^), anti-p-ULK1 (Ser^556^), anti-IRF3, anti-p-IRF3 (Ser^396^), anti-STING, anti-p-STING (Ser^366^), anti-GAPDH, anti-S6, anti-PERK, anti-IFIT1, anti-CYLD, anti-TAX1BP1, anti-PIAS1 and anti-TLR7 all from Cell Signaling Technology; rabbit polyclonal antibodies anti-eIF2α, anti-TBK1, p-S6 (Ser^235/236^) and 4E-BP1 from Cell Signaling Technology; anti-p62 (from Invitrogen); anti-SGK3 and anti-GADD34, both from Proteintech. HRP-conjugated secondary antibodies were purchased from Jackson ImmunoResearch (donkey anti-mouse) and from Cell Signaling Technology (goat anti-rabbit).

### Quantification of cytokines by ELISA

Human TNF-α was quantified in cell supernatants using a commercially available ELISA kit from PeproTech (900-TM25) and the appropriate commercial buffers (TMB ELISA Buffer Kit, 900-T00, PeproTech). LumiKin Xpress hIFN-β 2.0 (InvivoGen) was used to quantify the levels of human IFN-β. The assays were performed according to the manufacturer’s instructions.

### Confocal microscopy

For immunostaining, 5 × 10^4^ cells were centrifuged, resuspended in 50 µL of RPMI 1640 medium and added to Alcian blue-pretreated coverslips. After incubating 20 min at 37°C in 5% CO_2_, samples were fixed in 4% paraformaldehyde (PFA) (Alfa Aesar) for 10 min at RT and then washed three times in PBS. Cells were permeabilized and blocked with staining buffer (10 mM glycine, 5% FBS, 0.05% saponin, 1xPBS) for 10 min, before an overnight incubation at 4°C with primary antibodies diluted in staining buffer, in a wet chamber. Following three washes in staining buffer, staining with fluorescent secondary antibody (for unconjugated primary antibody) was performed for 1 h at RT, in a wet chamber protected from light. Finally, coverslips were washed three times in PBS, once in milli-Q water and were mounted in ProLong Gold Antifade Mountant (Invitrogen). Images were visualized in a Zeiss LSM 880 confocal microscope using a 63×/1.4 NA oil-immersion objective, being processed with ZEN 3.3 (blue edition) software (ZEISS). The antibodies used were rabbit monoclonal anti-EEA1 (Invitrogen), mouse monoclonal anti-LAMP1-BV421 (BioLegend) and goat polyclonal anti-rabbit Alexa Fluor 568-conjugated secondary antibody (Invitrogen).

### Mitochondrial ROS and cell viability measurement by flow cytometry

For detection of mitochondrial ROS, 5 × 10^5^ cells were centrifuged at 300 x g for 6 min and resuspended in 100 μl of RPMI 1640 medium. Samples were then stained with MitoSOX (Invitrogen) at 2.5 μM for 30 min at 37°C and 5% CO_2_. A wash in PBS was performed prior to acquisition on a BD Accuri C6 Flow Cytometer. To determine cell viability, cells were washed in PBS and mixed with propidium iodide (Sigma-Aldrich) at 7.5 µg/ml before acquisition. Data were analyzed with FlowJo software (Tree Star).

### Protein synthesis measurement by flow cytometry

To monitor protein synthesis, puromycin labeling was performed as previously described (Schmidt et al., 2009). At the end of the different treatments, cells were incubated with puromycin (Sigma-Aldrich) at 1 µg/ml for 15 min at 37°C and 5% CO_2_. Sodium arsenite (500 µM) (Sigma-Aldrich) was added 15 min before puromycin to be used as negative control for its incorporation. Samples were washed in PBS and incubated 30 min at 4°C in the dark with LIVE/DEAD fixable green dead cell stain (Invitrogen). Following a wash in PBS, cells were fixed in 4% PFA for 15 min at RT and were washed 3 times in PBS. A 15-min incubation in FACS solution (0.5% BSA, 0.01% sodium azide, 0.1% saponin, PBS) at 4°C was performed for cell permeabilization, before staining on ice in the dark for 1 h with mouse monoclonal anti-Puromycin-AF647 (Merck Millipore), diluted in the same solution. After 4 washes in FACS solution, cells were washed and resuspended in PBS and analyzed on a BD Accuri C6 Flow Cytometer. FlowJo software (Tree Star) was used to analyze samples.

### Statistics

GraphPad Prism Software was used for statistical analysis. Data representing multiple experiments are displayed as mean ± standard deviation of the mean (SD). Statistical significance was determined by paired Student’s t-test, or by one-way ANOVA followed by Dunnett’s, Tukey’s or Šidák’s multiple comparisons test, as indicated. *P < 0.05, **P <0.01, ***P < 0.001, ****P < 0.0001.

## Acknowledgements

Some figures were partly generated using Servier Medical Art, provided by Servier, licensed under a Creative Commons Attribution 3.0 unported license.

## Author contributions

P.A., M.D.M., F.L.P., D.B., A.M., B.H.F., M.R., C.R., D.C., I.F.D, M.N., R.J.A and B.N. performed research. B.S., P.P., C.R.A. and E.G. designed research and analyzed data. P.A., P.P., C.R.A. and E.G wrote the paper.

## Data Availability

This study includes no data deposited in external repositories.

## Expanded View Figure legends

**Figure EV1. Graphical abstracts.** Scheme of VPS34 functions (**A**), mTORC1 signaling (**B**), RLR and cGAS-STING signaling pathways (**C**) and unfolded protein response (**D**). (**E**) Summary of VPS34-IN1 impact on pDCs.

**Figure EV2. VPS34-IN1 acts efficiently in TLR7-activated CAL-1 cells.** (**A**) LC3B lipidation was analyzed by determining the LC3-II/LC3-I ratio. (**B**) CAL-1 cells were stained with MitoSOX and analyzed by flow cytometry. The mean fluorescence intensity was compared to untreated cells (ctr). (**C**) Immunofluorescence confocal microscopy of CAL-1 cells, showing EEA1 (red) and LAMP1 (blue) distribution. Scale bar, 5 μm or 1 μm in magnified areas. (**A**, **B**) Bars show the mean ± SD and each dot shows data from one experiment. ns, non-significant results; **P < 0.01, ***P < 0.001, by one-way ANOVA with Šidák post hoc test (A) and by one-way ANOVA with Tukey post hoc test (B).

**Figure EV3. Characterization of *ATG5^−/−^* and *UVRAG^−/−^* CAL-1 cell lines.** (**A, B**) Alignment of partial *ATG5* (A) and *UVRAG* (B) DNA sequences obtained in *ATG5^−/−^* and *UVRAG^−/−^* CAL-1 clones. Corresponding amino acid (a.a.) sequences are shown. Frameshift mutations in knockout cell lines are indicated; asterisk represents a premature stop codon in the coding sequence. (**C**) Representative immunoblot of LC3B (LC3-II – lipidated form; LC3-I – nonlipidated form) and ATG5 in wild type (WT) and *ATG5^−/−^* CAL-1 cells treated with bafilomycin. *ATG5^−/−^* clone (cl.) number 2 was selected for the experiments of this work. (**D**) Immunofluorescence confocal microscopy of untreated WT and *UVRAG^−/−^* CAL-1 cells showing EEA1 (red) and LAMP1 (blue) distribution. Scale bar, 5 μm or 1 μm in magnified areas. (**E**) Immunoblot detection of UVRAG in WT and *UVRAG^−/−^* CAL-1 cells.

**Figure EV4. VPS34-IN1 inhibits glycolysis in TLR7-stimulated CAL-1.** (**A**) Levels of glucose (left) and lactate (middle) were analyzed by NMR in cell culture supernatants of CL307-stimulated CAL-1 cells with or without VPS34-IN1; acellular medium was used as reference. Intracellular lactate levels (right) were monitored as well. Bars show the mean ± SD and each dot shows data from one experiment. ns, non-significant results; *P < 0.05, **P < 0.01, ***P < 0.001, ****P < 0.0001, by one-way ANOVA with Tukey post hoc test.

**Figure EV5. Mutations in DNA sequences of *MAVS^−/−^* and *STING^−/−^* CAL-1 clones.** (**A, B**) Alignment of partial *MAVS* and *STING1* DNA sequences obtained in *MAVS^−/−^* and *STING1^−/−^* CAL-1 cell lines. Corresponding amino acid (a.a.) sequences are shown. Frameshift mutations are indicated; asterisk represents a premature stop codon in the coding sequence. Immunoblot detection of MAVS and STING in wild type (WT), *MAVS^−/−^* and *STING1^−/−^* CAL-1 cells.

**Figure EV6. Characterization of *SQSTM1^−/−^* and *ULK1^−/−^* CAL-1 clones.** (**A, B**) Alignment of partial *SQSTM1* and *ULK1* DNA sequence obtained in *SQSTM1^−/−^* and *ULK1^−/−^* CAL-1 cells. Amino acid (a.a.) sequences are shown. Frameshift mutation is indicated; asterisk represents a premature stop codon. (**A**) Immunoblot detection of p62 in WT and *SQSTM1^−/−^* CAL-1 cells. (**C**) Immunoblot of LC3B and p-ULK1 (Ser^556^) in WT and *ULK1^−/−^* cells subjected to starvation (EBSS for 6 h), in the presence of bafilomycin. Quantification of LC3-II/LC3-I ratio is shown on the right. *ULK1^−/−^* clone number 11 was used in the experiments of this work. (**D**, **E**) WT and *ULK1^−/−^* CAL-1 cells were activated with 2’3’-cGAMP with or without VPS34-IN1. (**D**) Levels of total and phosphorylated forms of STING (Ser^366^), TBK1 (Ser^172^) and IRF3 (Ser^396^) and (**E**) levels of *IFNB1* mRNA were monitored after treatment with cGAMP and VPS34-IN1. (**C**-**E**) Bars show the mean ± SD and each dot shows data from one experiment. ns, non-significant results; *P < 0.05, ***P < 0.001, by paired Student’s t test.

## Appendix

**Appendix Table S1:** List of reagents, kits and cell models.

**Appendix Table S2:** List of gRNA genomic target sequences for CRISPR/Cas9 knockout

**Appendix Table S3:** List of primers

**Appendix Table S4:** List of antibodies.

**Appendix Supplementary methods**

